## Supplementary Information for "Distinguishing excess mutations and increased cell death based on variant allele frequencies"

#### S1 Implementation details

The synthetic sample generating code and the parameter estimation code can be accessed at `https://github.com/tg433/evolgenom`. Both are written in the Julia language.

#### S2 Synthetic data

For generating synthetic data, trees having 10000 leaves were used, generated by the ELynx software suite, with birth rate 1 and death rate, equaling the death-to-birth ratio, set to values $1 - \delta = 10^0, 10^{-1}, 10^{-2}, 10^{-3}, 10^{-4}$. For generating mutations, the corresponding mutation rates were $\mu = 6.5 \cdot 10^{-8}, 1.0 \cdot 10^{-8}, 2.5 \cdot 10^{-9}, 8.5 \cdot 10^{-10}, 5.0 \cdot 10^{-10}$. For the same death-to-birth ratio value, different datasets were generated using different trees. The chosen death-to-birth ratio-mutation rate value pairs resulted in approximately $5 \cdot 10^4$ observed mutations, in each case. No contamination by healthy cells and no clonal mutations were simulated. Sequencing depth was set to 100. The number of DNA sites was 3088286401, ploidy was 2.
To estimate the death-to-birth ratio and the mutation rate, 10000 trees were used at each of a selected set of death-to-birth ratios. The death-to-birth ratio points were equidistant on log-scale, the number of points within a decade being varied according to the distance from the true death-to-birth ratio value: 6-6 grid points in the 2 decades next to the true death-to-birth ratio value, 4 points in the second closest decades, 2 points for decades farther away. The interval of death-to-birth ratio was between $1-t = [10^{-6}, 10^0]$. Average loglikelihood values were calculated for those selected death-to-birth ratios; between them cubic

spline interpolation was utilized. The trees had 10000 leaves, similarly to the ones used for generating data, which were not used to estimate the parameters. Error rate and the number of clonal mutations were fixed at 0, contamination was also set to 0.

Extending Fig. 1. in the main text, Fig. S1 shows the illustrates the effect on the VAF of changing the mutation rate over three orders of magnitude for the two different death-to-birth ratios shown in Fig. 1.

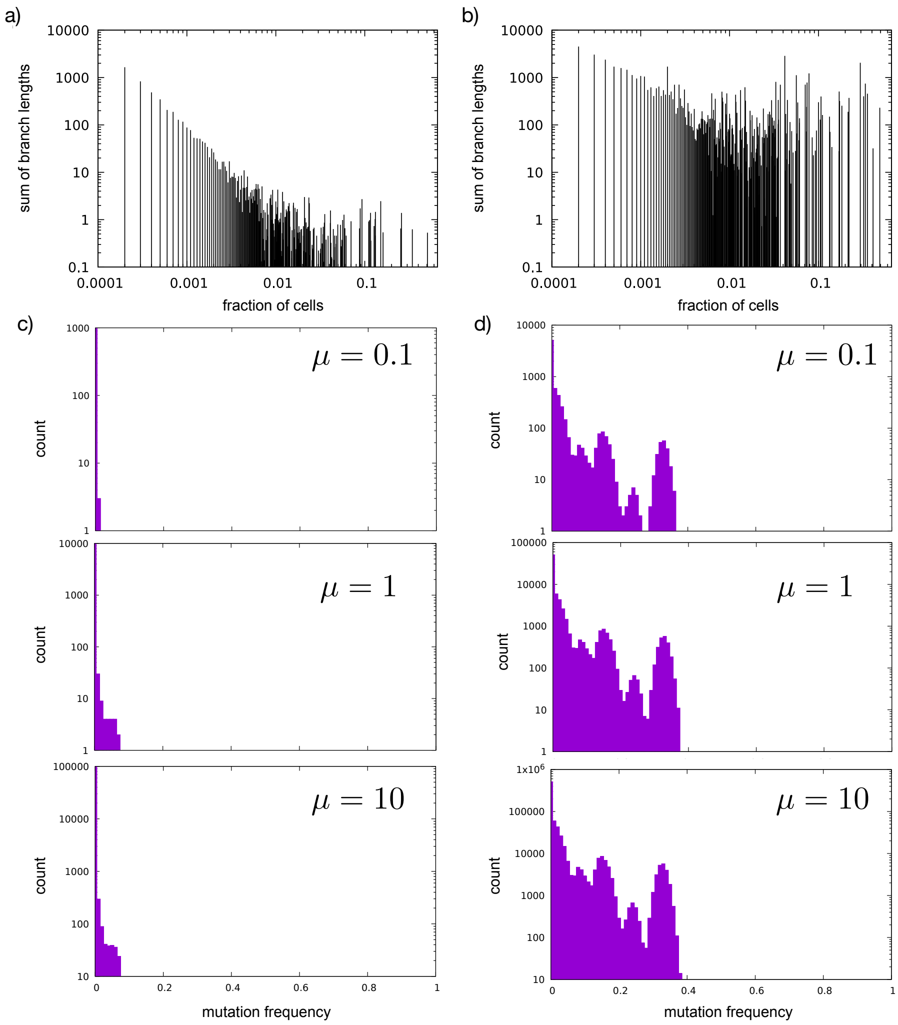


Figure S1: **VAF spectra for varying mutation rates and death-to-birth ratios.** Branch length distributions for a) low death rate and b) high death rate and corresponding VAFs, respectively c) and d), with the mutation rate varying over two orders of magnitude.

For Fig. 3. in the main text, datasets of $5 \cdot 10^5$ and $5 \cdot 10^3$ mutations were generated by simply adjusting the mutation rates by 10 and 0.1.

For Fig. 4. in the main text, different tree sizes were matched with different mutation rates, to maintain the number of observed mutations. For death-to-birth ratios $1-\delta = 10^0, 10^{-1}, 10^{-2}, 10^{-3}, 10^{-4}$ the mutation rates were $3.3 \cdot 10^{-7}, 1.0 \cdot 10^{-7}, 6.1 \cdot 10^{-8}, 4.5 \cdot 10^{-8}, 5.2 \cdot 10^{-8}$ (tree size 100), $1.1 \cdot 10^{-7}, 2.5 \cdot 10^{-8}, 8.4 \cdot 10^{-9}, 4.9 \cdot 10^{-9}, 3.0 \cdot 10^{-9}$ (tree size 1000), $1.6 \cdot 10^{-10}, 2.6 \cdot 10^{-11}, 6.1 \cdot 10^{-12}, 2.1 \cdot 10^{-12}, 1.3 \cdot 10^{-12}$ (tree size 100000). For tree size $10^5$, 100 trees was used to calculate the loglikelihood at each death-to-birth ratio value, due to the elevated computational requirements.

For Fig. 5. in the main text, synthetic data with simulated sequencing errors was generated similarly as before, simply using the error rates stated in the main text. The errors were generated by an independent stream of random numbers, so the underlying, error-free data are the same for all error rates.

### S3 Empirical data

Sequence data was obtained from GenBank with the project accession number of PRJNA339672 (`https://www.ncbi.nlm.nih.gov/bioproject/PRJNA339672`).

Preprocessing of data was done by following the steps described in [2], using the software tools provided by the authors. The local realignment step was omitted, as the current version of the proposed GATK software lacks the required tool, and VarScan 2, utilized later in the pipeline, does not need realignment.

Death-to-birth ratio values at which the loglikelihood values were evaluated are distributed evenly on log-scale, 4 points within a decade, between $1-\delta = 10^{-5}$-$10^0$. Between $1-\delta = 10^{-6}$-$10^{-5}$, 2 points are used.

Fig. S2 shows the loglikelihood-error rate plots of the empirical data. Different data points correspond to loglikelihood calculations using fixed error rate values. The optimal error rate is clearly very close to $\varepsilon = 10^{-7}$.

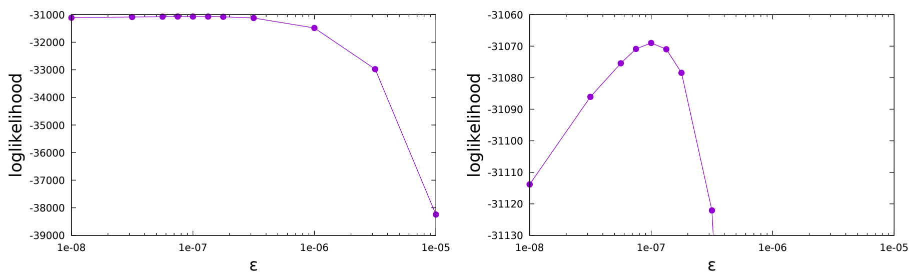


Figure S2: Loglikelihood-error rate curve of the empirical data. The right panel shows the peak with a narrow loglikelihood range.

Fig. S3 shows the variation of the death-to-birth ratio and the mutation rate according to the error rate. Both quantities vary only slightly even when the error rate is one order of magnitude away from its optimal value.

Fig. S4 shows the loglikehood as function of the death-to-birth ratio and mutation rate, for error rate $\varepsilon = 10^{-7}$.

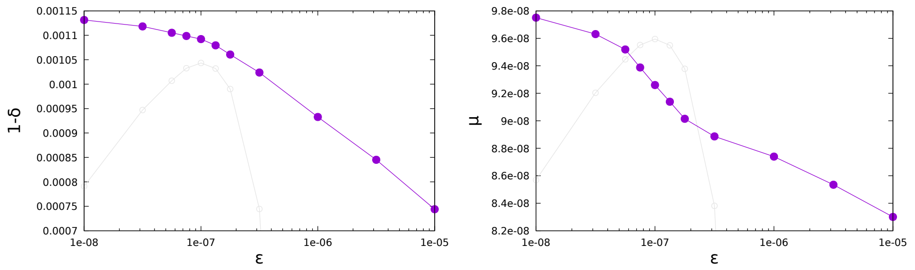


Figure S3: Death-to-birth ratio-error rate and mutation rate-error rate curves of the empirical data. The loglikelihood curve is shown in light gray.

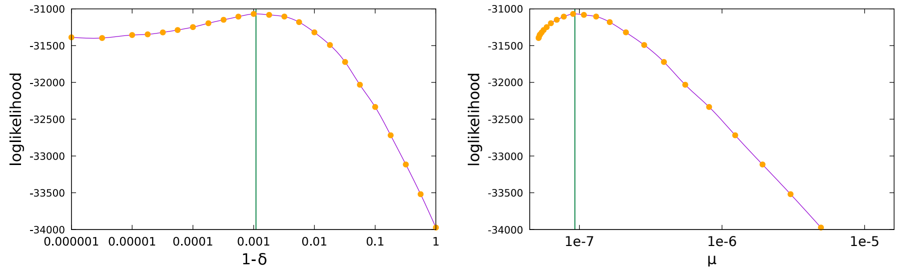


Figure S4: Loglikelihood-death-to-birth ratio and loglikelihood-mutation rate curves of the HCC data. Interpolation between data points is by cubic splines. Green line highlights the maximum.

### S4 Supplementary figures

Fig. S5 shows that resolving large death-to-birth ratios requires trees with more leaves.
Fig. S6 shows the estimated mutation rates in the presence of sequencing errors.

Fig. S7 shows the estimated mutation rates in the presence of sequencing errors.

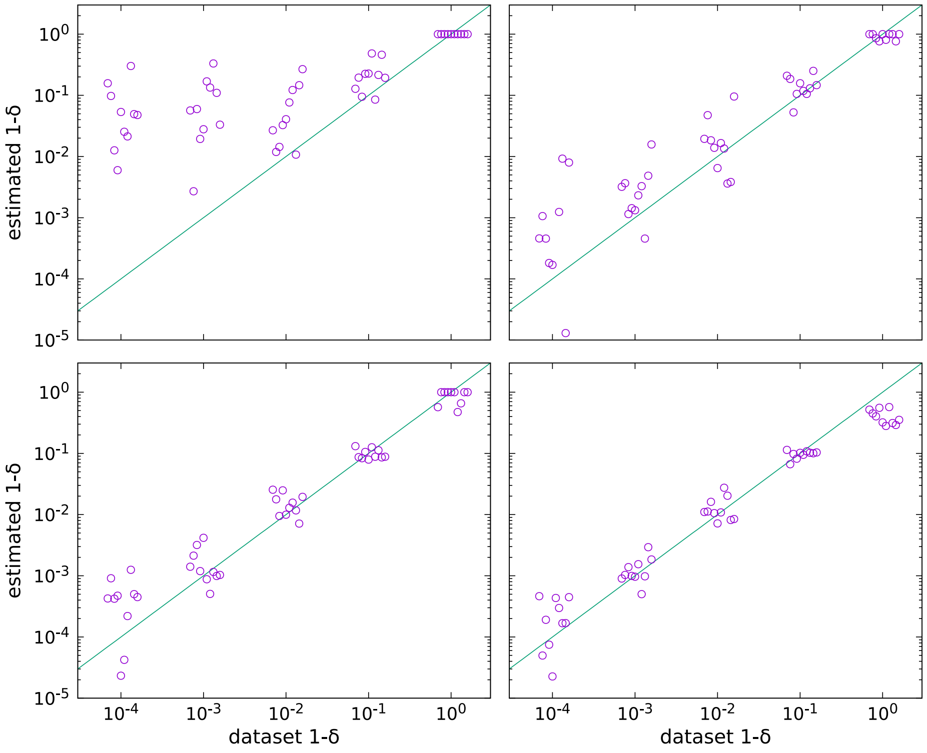


Figure S5: **The effect of tree sizes on the estimations.** Estimated death-to-birth ratios for fitted trees with 100 (top left), 1000 (top right), 10000 (bottom left), and 100000 (bottom right) leaves. Sizes of the sample generating trees are the same as those of the fitting trees.

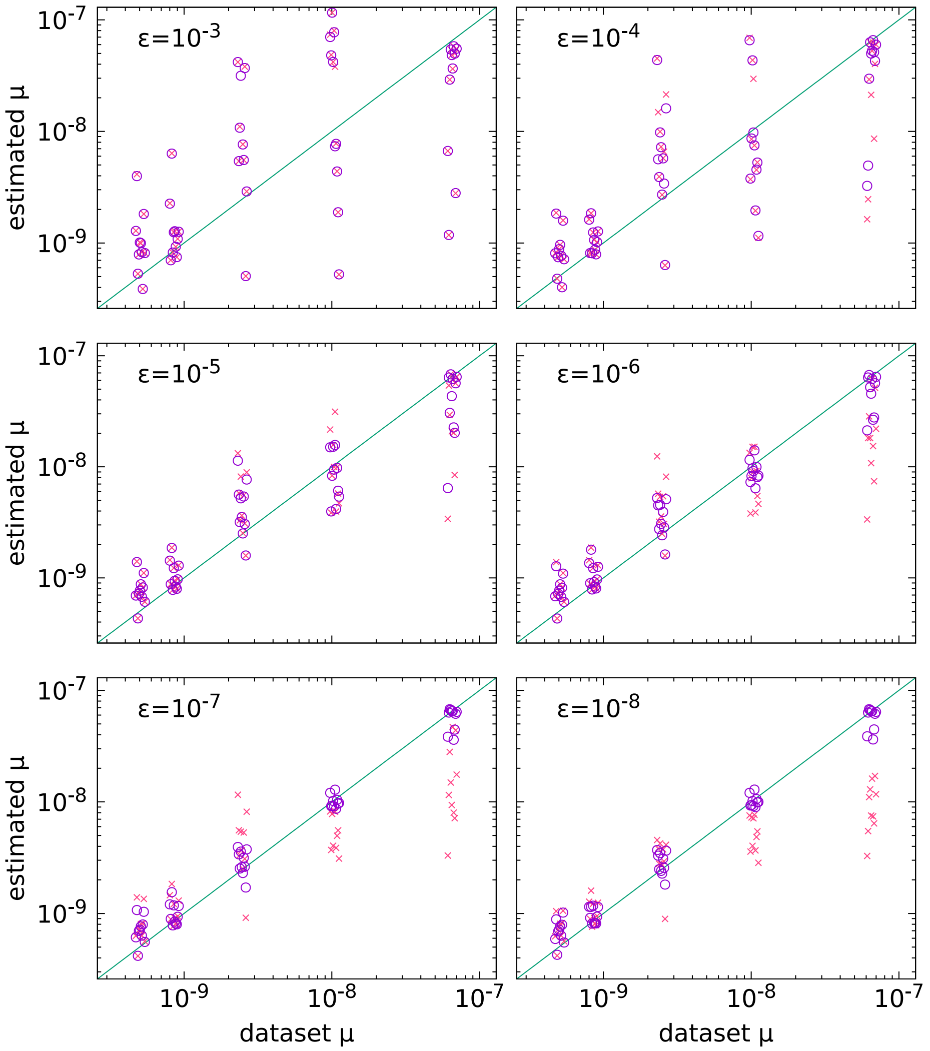


Figure S6: **The effect of the error rate on the estimated mutation rate.** Sequencing error rates are $\varepsilon = 10^{-3}$ (top left), $10^{-4}$ (top right), $10^{-5}$ (middle left), $10^{-6}$ (middle right), $10^{-7}$ (bottom left), $10^{-8}$ (bottom right). $10^4$ trees with $10^4$ leaves were used for fitting. Horizontal coordinates are slightly dispersed for clarity. Open circles are results corresponding to error rates fixed to their true values, crosses correspond to error rates estimated by the parameter fit. Each open circle-cross pair corresponding to the same dataset is vertically aligned.

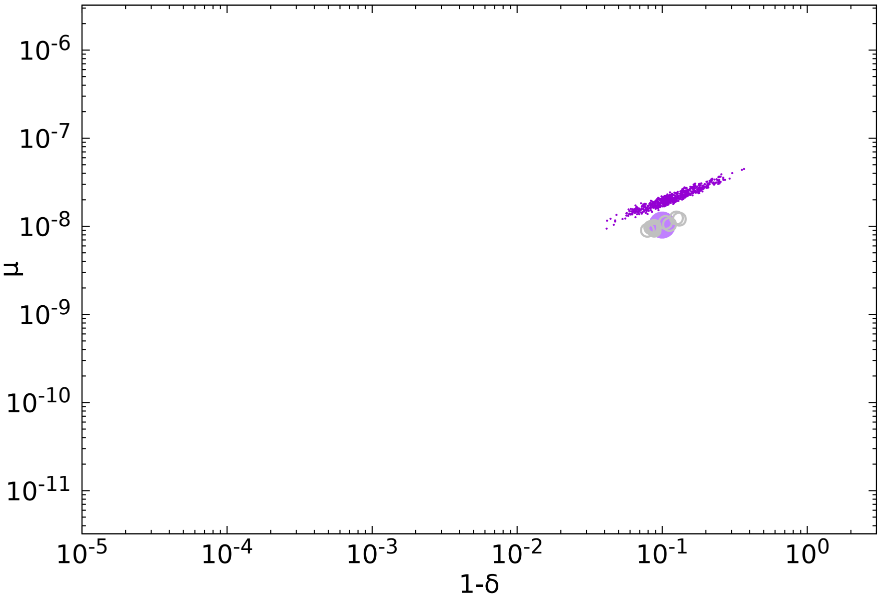


Figure S7: **The effect of the error rate on the estimated mutation rate.** In 500 replicate experiments we introduced a "driver" mutation that increases $1-\delta$ by 30%. The large purple point indicates the parameter values used to simulate the data, gray circles are inferences for data without driver mutations from Fig. 3, and purple datapoints are inferences for data including drivers. A small, but systematic increase is apparent in the inferred values of both the mutation rate and the death-to-birth ratio compared to "wild type" values resulting from the subtree with an increased death-to-birth ratio driver lineage.
